# VISTA: Virtual ImmunoSTAining for pancreatic disease quantification in murine cohorts

**DOI:** 10.1101/2020.04.01.020842

**Authors:** Luke Ternes, Ge Huang, Christian Lanciault, Guillaume Thibault, Rachelle Riggers, Joe W. Gray, John Muschler, Young Hwan Chang

## Abstract

Mechanistic studies of pancreatic disease progression using animal models require objective and quantifiable assessment of tissue changes among animal cohorts. Disease state quantification, however, relies heavily on tissue immunostaining, which can be expensive, labor- and time-intensive, and all too often produces uneven staining that is prone to variable interpretation between experts and inaccurate quantification. Here we develop a fully automated semantic segmentation tool using deep learning for the rapid and objective quantification of histologic features using hematoxylin and eosin (H&E) stained pancreatic tissue sections acquired from murine pancreatic cancer models. The tool was successfully trained to segment and quantify multiple histopathologic features of pancreatic pre-cancer evolution, including normal acinar structures, the ductal phenotype of acinar-to ductal metaplasia (ADM), dysplasia, and the expanding stromal compartment. Disease quantifications produced by our computational tool were highly correlated to the results obtained by immunostaining markers of normal and diseased tissue (DAPI, amylase, and cytokeratins; correlation score= 0.9, 0.95, and 0.91, respectively) and were able to accurately reproduce immunostain patterns. Moreover, our tool was able to distinguish ADM from dysplasia, which are not reliably distinguished by immunostaining, and avoid the pitfalls of uneven or poor-quality staining. Using this tool, we quantified the changes in histologic feature abundance for murine cohorts with oncogenic Kras-driven disease at 2 months and 5 months of age (n=12, n=13). The calculated changes in histologic feature abundance were consistent with biological expectations, showing an expansion of the stromal compartment, a reduction of normal acinar tissue, and an increase in both ADM and dysplasia as disease progresses (p= 2e-6, 6e-7, 4e-4, and 3e-5, respectively). These results demonstrate the tool’s efficacy for accurate and rapid quantification of multiple histologic features using an objective and automated platform. Our tool promises to rapidly accelerate and improve the quantification of altered pancreatic disease progression in animal studies.

## Introduction

Advances in deep learning technologies are creating opportunities for the rapid and objective assessment of both normal tissue and pathologic processes in biologic specimens. Computer-aided interrogation of medical imaging is being applied to accelerate and improve diagnosis in human patients [1, 2, 3, 4]. Similarly, deep learning technologies can greatly improve analyses in animal disease models which require the measurement of disease progression in large numbers of tissue samples resulting either from pharmacological or genetic manipulations. The extensive and growing use of murine models in disease studies creates a significant need for tissue assessment methods that are rapid, objective and quantifiable in order to permit statistically validated disease measurements among animal cohorts, free of technical variability and investigator bias.

The challenge of objective quantification of tissue changes among animal cohorts is significant. Evaluation of tissue by either histochemical stains or antigen-specific immunohistochemistry offers distinct and sometimes overlapping information, but both have limitations. Hematoxylin and eosin (H&E) staining is a rapid, reliable and inexpensive method; however, lack of molecular specificity and requirement for manual segmentation have, thus far, limited its use for extraction of quantifiable data. Consequently, disease assessments by H&E staining are typically qualitative and vulnerable to inter-observer variation and bias [5,6,7]. Immunohistochemical stains offer a degree of specificity, but immunostaining can be labor- and time-intensive, expensive and results are often challenging to objectively quantify over broad tissue regions. In addition, tissue features of interest are not always cleanly distinguishable by immunostaining markers, and so tissue assessments can be limited by reliance on the molecular specificity of antibodies.

Using murine models of pancreatic cancer progression and pancreatitis, we are working to develop and validate deep learning approaches that enable the rapid, reliable, and automated quantification of disease progression over large tissue areas, solely based on H&E staining. Murine models of pancreatic cancer were chosen as they have proven useful for mechanistic investigations of pancreatic cancer progression, modeling well the human disease both genetically and phenotypically, particularly during the evolution of pre-cancerous lesions [8, 9]. The murine models have produced an explosion of studies including pre-clinical drug tests and evaluation of additional genetic perturbations that expose tumor-suppressing and tumor-promoting disease modifiers [10-12].

The early stages of pancreatic cancer evolution are well described in the mouse models [8, 9]. The normal pancreas consists predominantly of acinar and ductal epithelial cells forming the exocrine compartment, along with islet cells of the endocrine compartment, vasculature and the varied fibroblasts of the stromal compartment. The earliest stages of oncogene-induced pre-cancer evolution are marked by an expansion of ductal cells or by the conversion of the acinar cells to a ductal phenotype in an adaptive process known as acinar-to ductal metaplasia (ADM) [13]. ADM is also characteristic of acute and chronic pancreatitis, inflammatory conditions that can predispose to cancer [13]. The next stage in cancer evolution is the development of low-grade dysplasia, also referred to as pancreatic intraepithelial neoplasias (PanINs 1 and 2). Low-grade dysplasia is a pre-invasive neoplasia that can evolve to high-grade dysplasia (PanIN 3) and then progress to invasive pancreatic ductal adenocarcinoma (PDAC) [14]. Both ADM and dysplasia are accompanied by a prominent stromal reaction and immune cell infiltrate [13]. The stages of ADM and dysplasia evolution are believed to encompass a long phase of pre-cancer evolution that is a valuable window for early intervention [14].

Here we describe the model training workflow and application of deep learning on H&E stained samples of murine pre-cancerous lesions, segmenting the normal acini, the ductal phenotype of ADM, and dysplasia. With the rapid growth of computer vision, more specifically deep learning, novel image analysis architectures have been developed for accessing image information that is not readily observed through traditional methods. Several research groups have worked towards inter-modality image translation and have developed tools that attempt to convert medical images such as H&E stained tissue and brightfield microscopy to more detailed ones such as fluorescent immunostains [15,16,17,18]. The target of such models has been the direct translation of stain intensities for the purpose of constructing entirely new images. Our developed tool seeks to go further, predicting binarized masks of positive staining area and augment immunostaining by segmenting key histologic features that current stains cannot reliably differentiate.

Results presented here demonstrate a well validated segmentation tool that can automatically, rapidly, and objectively quantify pancreatic tissue and disease progression in mice, relying solely on easily replicated and low-cost H&E staining of whole pancreas tissue sections, free of experimental variability and investigator bias. Our work provides a tool that is immediately applicable to the improvement and acceleration of pancreatic disease studies in animal cohorts, and provides workflows for similar tool development in other disease models. Moreover, the ease of use and availability allows for this tool to be a common thread for comparing different studies performed throughout the world.

## Results

Murine pancreatic pre-cancerous tissues were isolated from the *P48*^*+/Cre*^; *LSL-KRAS*^*G12D*^ mice (KC) which is a common mouse pancreatic cancer model that displays the early disease hallmarks of ADM, dysplasia, and desmoplasia, and can eventually develop invasive adenocarcinoma after more than one year of age [8]. Tissue sections from 3 whole pancreases were acquired from KC mice at 5 months for models training, and whole pancreas sections from an additional 25 mice were collected at 2 and 5 months of age (n=12, n=13) for validation and testing on an unseen dataset. All pancreas tissue sections were stained with H&E and the validation set was additionally stained by immunofluorescence for amylase (AMY), labeling normal acini, pan-keratin (panK), labeling primarily the oncogenic Kras-transformed epithelial population, and DAPI, labeling all nuclei.

In order to predict the immunofluorescent stain and histologic feature distributions, several UNet models [19] were trained using intensity normalized H&E image tiles paired with ground truth tiles created from three experts’ annotations using CYTOMINE [20]. Annotations were generated to designate normal acinar structures, the ductal phenotype of ADM, and dysplasia. Given expert annotations as ground truth, models were trained by optimizing the Binary Cross Entropy Loss, and following training, the Dice Coefficient was used to select the best models. These trained models were then validated quantitatively by correlating their predicted stain distributions to a set of stained and binarized fluorescent images where the intensity threshold is chosen by an expert (Figure 1). As observed in Table 1, the models implementing Reinhard Normalization [21] achieved better scores on average, relative to Vahadane [22] and Macenko [23] normalization methods. Furthermore, the models achieved the best scores when the normalization process was applied on intermediate overlapping crops rather than across the whole image.

**Table 1:**
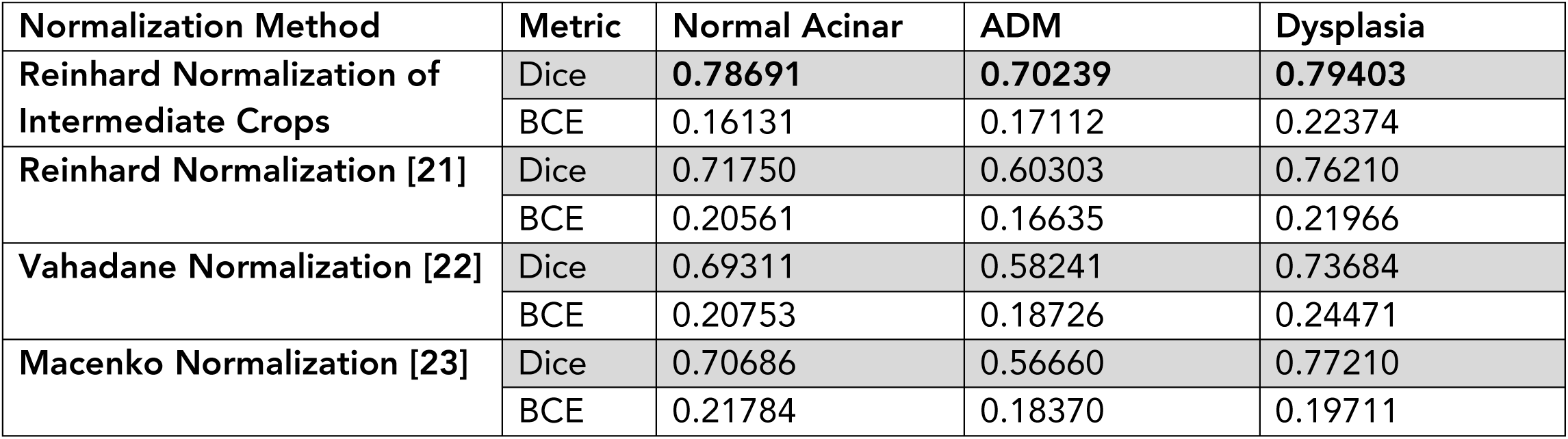
Evaluation of Model Performances.

| Normalization Method | Metric | Normal Acinar | ADM | Dysplasia |
| --- | --- | --- | --- | --- |
| Reinhard Normalization of Intermediate Crops | Dice | **0.78691** | **0.70239** | **0.79403** |
|  | BCE | 0.16131 | 0.17112 | 0.22374 |
| Reinhard Normalization [21] | Dice | 0.71750 | 0.60303 | 0.76210 |
|  | BCE | 0.20561 | 0.16635 | 0.21966 |
| Vahadane Normalization [22] | Dice | 0.69311 | 0.58241 | 0.73684 |
|  | BCE | 0.20753 | 0.18726 | 0.24471 |
| Macenko Normalization [23] | Dice | 0.70686 | 0.56660 | 0.77210 |
|  | BCE | 0.21784 | 0.18370 | 0.19711 |

**Table 2:** Number of Training Annotations:

|  | Number of Annotations |  |  |
| --- | --- | --- | --- |
|  | Normal Acinar | ADM | Dysplasia |
| Image 1 | 119 | 1722 | 1659 |
| Image 2 | 1342 | 597 | 70 |
| Image 3 | 463 | 263 | 3 |
| Total | 1924 | 2582 | 1732 |

**Figure 1:**
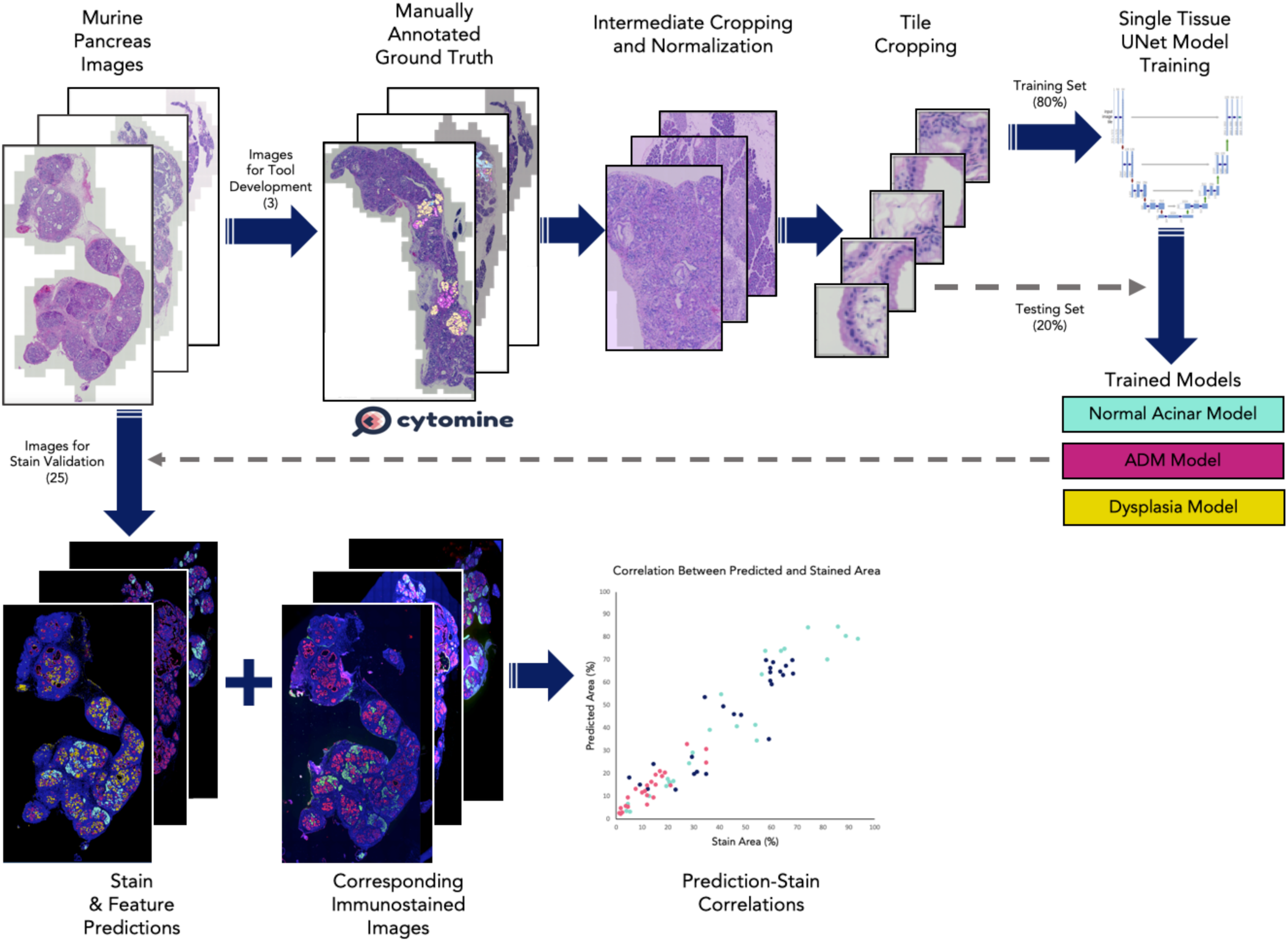
Experiment workflow. A subset of murine pancreas H&E images were annotated by three experts in Cytomine [20]. The images and their annotations were cropped and normalized at intermediate intervals, and these intermediate crops were then tiled into images that can be fed into a UNet architecture [19]. 80% of tiles were used for training and 20% were used for testing. A model was trained for each histologic feature label. The best models were chosen and used to predict stain and feature distributions on unseen H&E images. These predictions were then correlated with the stained image counterparts to determine model accuracy.

The models were trained using 80% of the training data, and 20% was held out for cross-validation to evaluate the models’ performance with unbiased data. The best models yielded Dice Coefficients of ∼0.79, ∼0.70, and ∼0.79 on the hold-out set for normal acinar tissue, ADM, and dysplastic features, respectively (Table 1). The segmentations match the expert annotations with a high degree of qualitative accuracy (Figure 2a). The models’ Dice scores are lower than expected from successful models is because the models actually refined approximations in the experts’ annotations leading to discrepancies between prediction and annotation (Figure 2b). Due to the limitations of the annotation method used, entire lesions, including empty lumina, were labeled as one type of tissue (i.e., ADM or dysplasia). The models, however, accurately differentiate between the tissue types within a lesion and avoid labeling lumina. Despite these results being biologically correct, they are different than the experts’ manual annotations, resulting in a negative impact on the measured Dice Coefficients.

**Figure 2.**
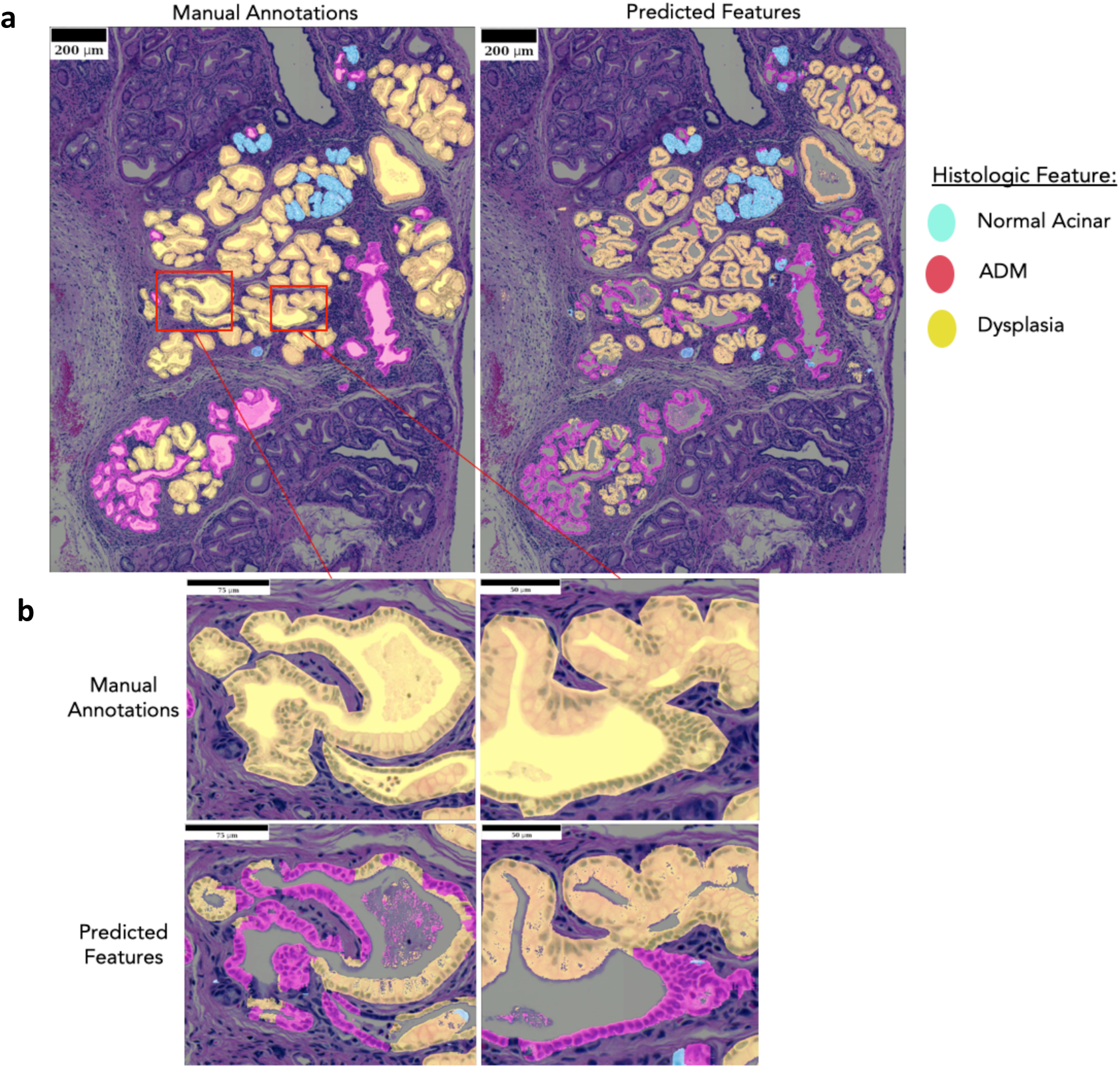
Predictions compared to annotations. a) Model Predictions closely align with the manually annotated ground truth regions that was used for training. b) Close inspection of the ducts shows consistent discrepancies regarding the lumen and split histologic features within single ducts. Manual annotations were made by circling whole ducts, but the models’ predictions are actually more reflective of biology, wherein, stain does not mark for the lumen. The Predictions can also distinguish histologic features differences that the manual annotations combined.

To test further the accuracy of the trained models, a comparison was made between quantified model predictions and quantified immuno-stained images that have been binarized. Quantification of the tissue area occupied by normal acinar cell and transformed pancreatic epithelial cells was achieved by immunostaining for amylase and pan-keratin, respectively, with DAPI staining of nuclei used to detect all cellular regions. The comparable calculation was then made using tool predictions on adjacent H&E stained tissue sections. For the tool prediction, ADM and dysplasia predictions were grouped into the panK stain because pan-keratin immunostaining does not distinguish ADM and neoplastic tissues. Because stain area is more biologically targeted than the rough annotations that incorporate empty lumens and mislabeled features, the models’ immunostain correlation scores are much more reflective of their overall accuracy. When the prediction masks are compared qualitatively to the stained images, the models are able to approximate the immunostain localization (Figure 3a). There are minor differences between the immunostained and the predicted segmentations, which reflects slight tissue variations between the adjacent, but separate, sections used for H&E staining and immunohistochemistry. Quantitatively, three models also have high correlations (Figure 3b) with the immunostained sections despite these sections (n= 25) being unseen during training. This validates that the models have been successfully trained and are capable of replicating known biological data.

**Figure 3.**
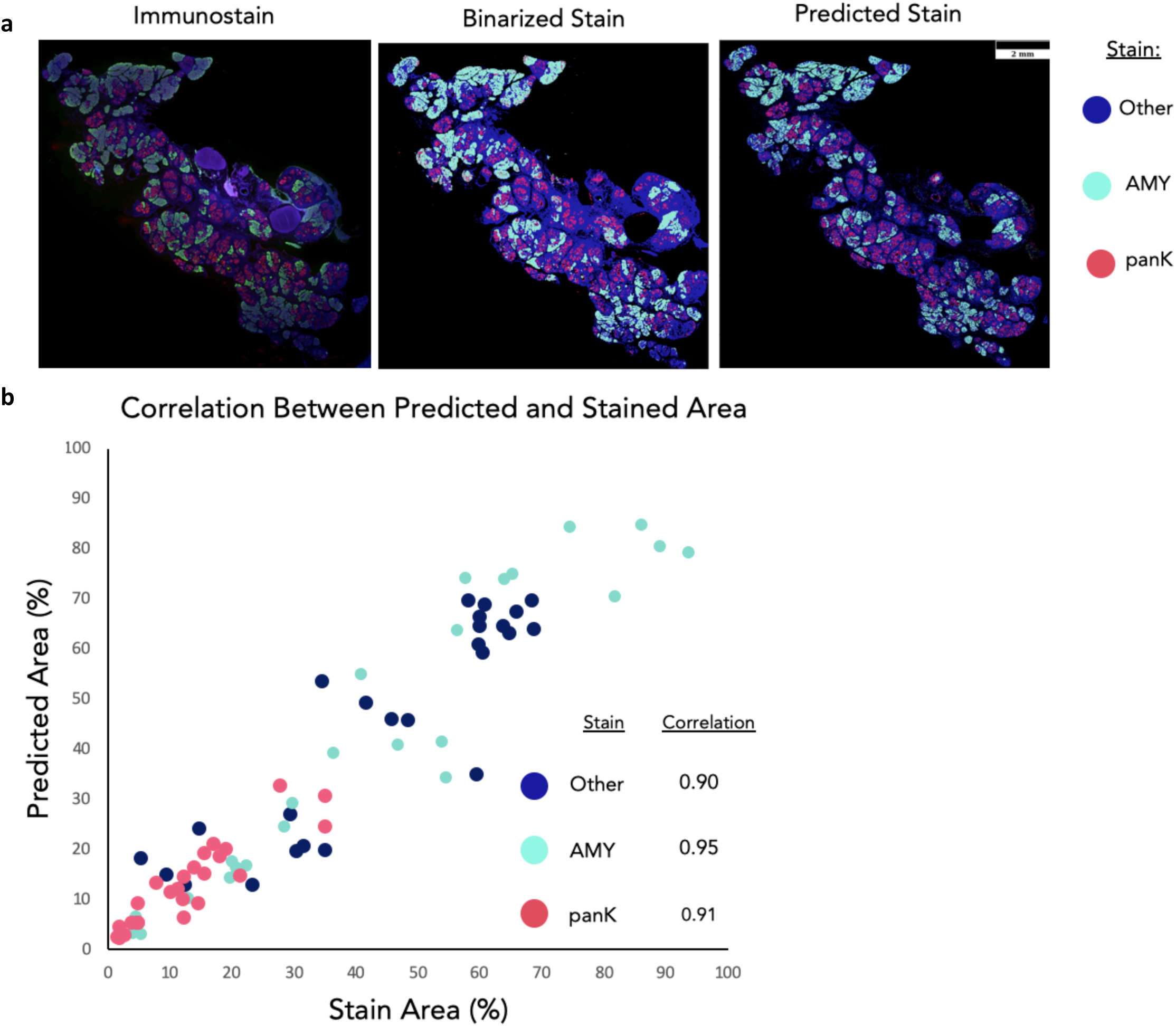
Comparing model predictions to stained tissue. a) Stain masks and predicted segmentation masks are qualitatively highly similar. Differences can be seen in the high-level architecture of the tissues, which is indicative of the fact that the predictions were made from serial sections to the stains. There are also dim regions of the stained image that are lost from the global thresholding technique. These regions are successfully captured by the models. “Other” stain is the DAPI stain minus regions overlapping with AMY and panK. b) Correlations were made by comparing the percent of area coverage for each stain mask. The high levels of correlation illustrate the models’ ability to replicate straining using only H&E images. These regions are successfully captured by the models. “Other” stain is the DAPI stain minus regions overlapping with AMY and panK.

Not only can these models replicate immunostaining data, they can extract more information than can be gained via immunostaining. In this dataset, the pan-keratin immunostain labels both metaplasia and dysplasia, restricting the disease features that can be segmented. The model predictions, however, can distinguish these features (Figure 4a). This allows for deeper and more nuanced quantification of disease progression than can be achieved by immunostaining alone. Because this process of prediction is deterministic, it is also a faster and less biased than manually annotating histologic features, and less expensive and less error-prone than immunostaining (Supplemental Figure 1).

**Figure 4.**
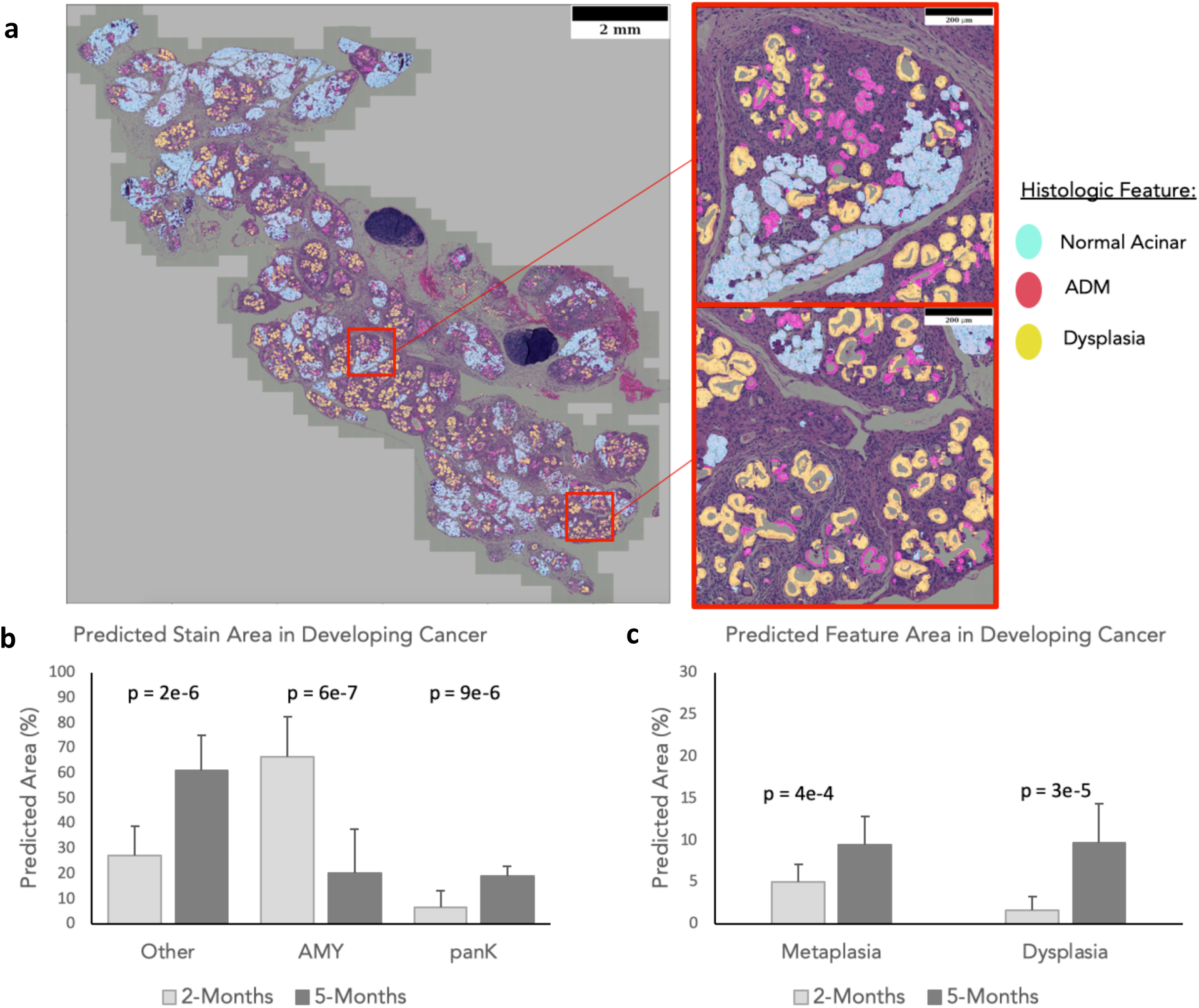
Discerning features beyond immunostaining. a) In test images the predicted histologic features visually align with what is expected from the H&E images. This shows the models’ utility in discerning novel information regarding ductal features that cannot be detected via staining. The models were used to predict the changes stain distributions b) and cancer histologic features c) in murine models with induced cancer. Predictions show significant changes in all stains and features between time points, and quantifies specific features that were not discernable in immunostaining alone.

Using the tissue sections from the unseen testing dataset isolated from KC mice at 2 and 5 months of age (n=12, n=13), the model was able to quantify tissue changes reflecting disease progression by predicting immunostain from H&E stain images (Figure 4b and c). The observed age-dependent transitions from normal acinar to ADM and dysplasia, and the increase in other tissue area (DAPI stained), is consistent with biological expectations, illustrating the practical, objective use of this tool to quantitatively assess pre-cancerous disease development.

To test the models’ robustness and generalizability, we evaluated images from pancreata with acute pancreatitis. Acute pancreatitis is characterized by prominent ADM and an inflammatory stromal response, but does not promote neoplastic lesions [13]. Acute pancreatitis was induced in mice by injection of the pro-inflammatory agent caerulein [13], then tissue sections exhibiting acute pancreatitis or normal pancreas (n=6, n=3 respectively) were analyzed by the model (Figure 5). Because annotations did not exist for these datasets, model prediction localizations were evaluated qualitatively. Despite not being trained to analyze the particular disease states of pancreatitis, the models were able to accurately label pancreatitis features (i.e. ADM) with minimal error, regardless of whether the ADM was sporadic or clustered within the tissue (Figure 5a). The model’s quantified tissue assessments show the significant presence of ADM by pixel area in the pancreatitis samples compared to normal tissues, which matches biological expectations. The near-absence of significant ADM and dysplasia in normal pancreas samples is also consistent with expectations, as is the near-absence of dysplasia in the pancreatitis samples (Figure 5b). The small quantities of ADM and dysplasia predictions in the normal tissues are errors introduced primarily by pixel level noise and are insignificant compared to the size of the samples. Within this dataset we do not see large heterogeneity in the histologic features across disease states, and as a result the model performs consistently across all disease states shown.

**Figure 5.**
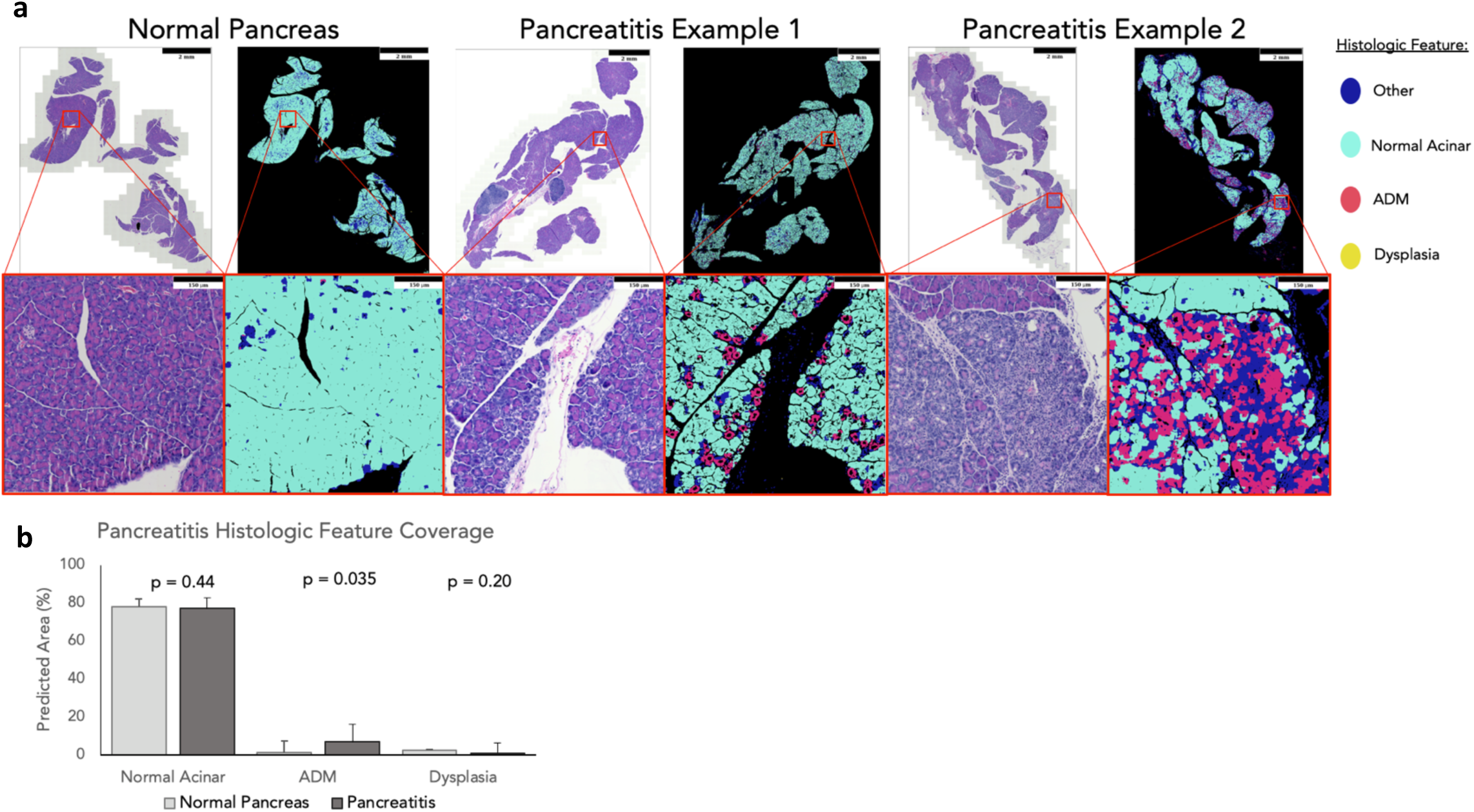
Predicting histologic features in pancreatitis. The model predicted histologic features match what in expected in both normal and pancreatitis samples. a) Predicted images show that tissue is dominated by normal acinar with pockets of clear ADM localization. In normal tissue ADM and dysplasia are sparse predictions comprised primarily of arbitrary single pixels, and in pancreatitis this is true for just dysplasia. b) In normal tissues, ADM and dysplasia predictions are negligible, and in pancreatitis there is a significant spike in ADM coverage with negligible dysplasia. Erroneous predictions of ADM and dysplasia in these samples are primarily driven by noise.

## Discussion

The computational tool developed here is intended to augment and accelerate disease research performed in animal models by allowing for simple stain prediction and histologic feature labeling from H&E images without the need for expensive and time-consuming immunostaining and biased image interpretation. It can be used to both mark the localization of tissue features and quantitatively to measure the extent of disease based on multiple histologic features (Supplemental Figure 2). Such rapid and unbiased quantification of disease states in animal models is critical to enabling efficient and accurate disease assessments among large study cohorts, as well as provide a common method to compare finding across different studies. The ability of this tool to accurately predict histologic features among 25 unseen pancreatic pre-cancer samples from multiple time points and 9 unseen samples comprising other disease states demonstrates the robustness of the models when analyzing new datasets. The fact that the models generalize well, despite being trained with a relatively small dataset (Supplemental Figure 3 and Table 2), illustrates the effectiveness of this workflow for tool development. Using this workflow makes niche tool development plausible for small working groups that might have less access to the resources needed to produce large batches of annotated data.

There have been many efforts to recreate advanced staining images using more common input modalities [15,16,17,18], and although they are useful for visualizing potential stain and intensity distributions, the algorithms are limited to predicting staining patterns of existing markers. If the user wants to analyze specific biological features for which there is no specific stain; however, simple stain translation will not suffice. The tool created here, however, can create objective binary interpretations of H&E images that segment histologic features of developing pancreatic cancer for which there is no reliable conventional immunostain. Previous studies have attempted to use computer-aided analyses for duct detection in pancreatic cancer [24]; however, these do not cover the subtly different features of early disease hallmarks of ADM and dysplasia.

Although this tool enables easy, rapid, and accurate stain reconstruction and feature labeling in the early stage disease models employed here, there are several limitations to its predictive capacity. The most prominent source of error for the tool currently is the way it handles unlearned tissue types, such as lymph nodes, pancreatic islets, the desmoplastic stroma, and the occasional presence of neighboring gastrointestinal tissue. Lymph nodes and gastrointestinal tissue are highly irregular compared to the pancreatic features that were present in the training data, leading to completely arbitrary labeling of the unrecognized tissue areas. To overcome this, these regions can simply be cropped prior to analysis, as performed for our analyses. Islets comprise a small fraction of the pancreatic tissue area, and were labeled by the model as “other” (i.e. neither normal, ADM, or dysplasia), and therefore introduced only minor errors. In addition, the desmoplastic stroma is a prominent and histologically distinct feature of pancreatic disease that is currently unlearned and labeled as “other” tissue.

Greater limitations arise with the appearance of high-grade neoplasia and adenocarcinoma, both of which can adopt ductal structures more closely resembling ADM. It should also be noted that the tool currently labels all non-neoplastic ductal structures as ADM, whether they originate from acinar cells or from ductal cells, and this contributes some error for the quantification ADM of acinar origin. At this stage of the tool’s development, no label for fully developed adenocarcinoma features were used, so lesions that have progressed beyond high grade dysplasia would likely be mislabeled as either ADM or “other”. With future work, it should be possible to train models to identify these additional tissue features and predict them accurately alongside the existing models. The final limitation of the tools is its failure to make accurate predictions in areas of tissue folding or out of focus imaging, but these are obstacles for any image-based measurement tool (including human annotators) and are avoidable with good technique.

Further work is in progress to reduce error and allow for a broader range of tissue interrogations, including training the tool to recognize a greater diversity of cell types and tissue features such as islets of Langerhans, neural tissue, desmoplastic stroma, adenocarcinoma, and peripheral elements such as lymph nodes or gastrointestinal tissue. The model’s quantitative capabilities can also be applied to other disease states that share similar histologic features, such as pancreatitis. Continued development can yield a single comprehensive tool for predicting and labeling all histologic features in pancreatic tissue without the need for complex staining.

Despite the current limitations discussed above, the tool developed here demonstrates clear advantages and superiority to immunostaining for disease quantification in pancreatic pre-cancers. By relying on H&E staining alone, the data acquisition is not only faster and cheaper, but less vulnerable to variable and uneven staining across tissue sections. This consistency and stability of H&E staining eliminates a primary source of error and bias in feature quantification because of manual adjustments needed to threshold immunostained tissues; tissue immunostaining quality varies significantly within single tissue sections and among the many tissues acquired and stained from animal cohorts, typically stained on different days, months, and even years. This tool’s exploitation of H&E staining not only enables easy quantitative comparisons between tissues collected and stained across broad time periods, but also enables such comparisons among tissues collected and stained in different laboratories around the world. This unifying aspect will improve collaboration and cross-validation between experiments conducted by different groups.

Being computer driven, the tool easily quantifies whole pancreatic tissue sections, allowing greater volumes of data acquisitions and avoiding the selection of “representative” regions for quantification, which introduces further bias. Furthermore, as an automated, machine-driven measurement tool, potential investigator bias is excluded from the data quantification pipeline. Finally, and importantly, tool has been demonstrated to identify and segregate key histologic features which immunostaining methods cannot reliably distinguish (i.e. ADM and dysplasias), significantly extending the power of available tissue analytics. This genre of tool will certainly enhance, and conceivably fully replace immunostaining in many animal studies.

## Competing Interests statement

The authors declare no competing financial or non-financial interests.

## Code availability

The tool’s code for making predictions is provided on GitHub at the following link: https://github.com/GelatinFrogs/MicePan-Segmentation

Images needed to run the tool can be found in the following google drive: https://drive.google.com/drive/folders/1ipgkjPawkuoLqtLENjHvSVC7hcZKWbRJ?usp=sharing

## Acknowledgements

This work was supported in part by the National Cancer Institute (U54CA209988, U2CCA233280), a Brenden-Colson Center for Pancreatic Care’s pilot (Y.H.C), a Brenden Colson Center for Pancreatic Care Fellowship (J.L.M.), and a Brenden-Colson Center for Pancreatic Care Program Leader Discovery Funding Project (J.W.G). YHC acknowledges the OHSU Center for Spatial Systems Biomedicine and Biomedical Innovation Program Award from the Oregon Clinical & Translational Research Institute.

## Author Contributions

LT developed algorithms, wrote the code, analyzed the data, and interpreted the results. GH performed the experiments and analyzed the immunofluorescent datasets. JM conceived and designed the study and GT and YHC supervised and oversaw the computational development. JM and RR performed tissue annotations, and CL provided pathology instruction and qualitative assessment. LT drafted the manuscript and all the authors reviewed the manuscript. All authors critically revised the manuscript and provide intellectual content.

## Methods

### Dataset

Murine pancreatic tissues displaying a range of pre-cancerous lesions were isolated from the *P48*^*+/Cre*^; *LSL-KRAS*^*G12D*^ mice (KC) mouse pancreatic cancer model. This a widely used genetically engineered mouse model of oncogenic Kras-driven pancreatic adenocarcinoma that closely models the evolution of the human disease, displaying the early hallmarks of ADM, Dysplasia, and desmoplasia, and eventually invasive adenocarcinoma after more than one year of age [8]. Tissue sections from 3 whole pancreases were acquired from KC mice at 5 months for models training, and whole pancreas sections from an additional 25 mice were collected at 2 and 5 months of age (n=12, n=13) for validation and testing on an unseen dataset. Collected pancreases displayed abundant pre-cancerous lesions but were preceding the development of adenocarcinoma. All pancreas tissue sections were stained with H&E and the validation set was additionally stained by immunofluorescence for amylase, labeling normal acini, pan-keratin, labeling primarily the oncogenic Kras-transformed epithelial population, and DAPI, labeling all nuclei.

### H&E staining and Immunofluorescence

The pancreatic tissues were paraffin-embedded, sectioned at 5μm thickness, and H&E stained by standard protocols at the OHSU Histopathology Core. For immunofluorescence staining of amylase and pan-keratin, antigen retrieval was performed using Dako Target Retrieval Solution at pH 9 (Aligent: S236784-2) according to manufacturer’s instructions. Specimens were blocked with blocking buffer (1X PBS/5% normal serum/0.3% Triton™ X-100) for 1 hour at room temperature. The anti-amylase (Santa Cruz: sc-12821) and anti-pan-Cytokeratin (Santa Cruz: sc-15367) primary antibodies were incubated overnight at 4°C, then washed and incubated with secondary antibodies (Invitrogen: A10042 and A32814) for 1.5 hours at room temperature. Slides were covered by coverslips with DAPI’s Prolong® gold anti-fading agent (Invitrogen: P36931). Fluorescent images of amylase (A), pan-cytokeratin (B), and DAPI (C) staining were acquired using a Carl Zeiss Axioscan Z1 slide scanner at a resolution of 0.2 microns/pixel and converted to BigTiff format.

Immunofluorescence images were quantified using ImageJ software. The threshold tool was applied manually to select the amylase-, pan-cytokeratin, or DAPI-positive tissue regions. Lymph nodes were manually cropped and excluded.

### Expert Annotation

Annotations for pancreatic tissue features were constructed in Cytomine [20] by three trained experts, and affirmed by a pathologist. These annotations came from 5 regions across 3 images (Supplemental Figure 3) and included at total of 1924 normal acinar, 2582 ADMs, and 1732 Dysplasia (Table 2).

### Training Image Preparation

In order to make the images more amicable to training for the Deep Learning algorithms, they were trained with intensity normalization to make them appear more consistent with each other. To overcome differential staining across an H&E image, various normalization approaches were applied on intermediate sized (5000×5000 pixel) overlapping crops prior to tiling (512×512 pixel). Background intensities were also ignored from the normalization process to reduce drastic changes on edge regions, isolating only the areas of interest for normalization. Background area was selected by thresholding pixels where all RGB values were greater than 200. The best normalization method was shown to be Reinhard normalization [21] (Table 1), so it is used in the implementation of the models.

### UNet Training

A separate UNet model was trained for each annotated ductal tissue type (normal acinar, ADM, and Dysplasia) [19]. To make each model specific to its respective tissue type, each model’s training set was made to incorporate small portions of the other tissue types as negative controls. The training sets were made using 80% of the total relevant tissue tiles and ∼5-10% of the total of other tissue tiles. Tiles were augmented during training with flips, rotations, and shears to overcome the small dataset size. Training for all three models lasted for 50 epochs, used a batch size of 32 tiles and had a learning rate of 7e-4, implementing the Adam optimizer. Binary cross entropy was used as the loss function during training. Dice Coefficient was used following training to select the best models.

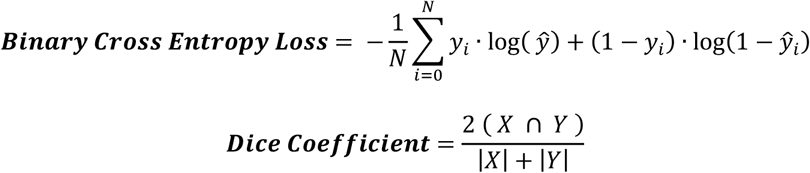

### Model Integration

Following model prediction, positive pixels for each model were calculated using the following thresholds:

Normal Acinar Threshold = 0.3, ADM Threshold = 0.5, Dysplasia Threshold = 0.7,

These thresholds were chosen based on the Receiver operating characteristic (ROC) curves (Supplemental Figure 4), and were manually adjusted to improve generalizability and reduce noise in the test images. Again, background white pixels were removed from prediction by ignoring all pixels where all RGB values were greater than 200. Total tissue (DAPI positive) region was also calculated by finding all pixels where RGB values were lower than 200. To combine all four tissue masks, normal acinar predictions override metaplasia and dysplasia predictions; metaplasia predictions override dysplasia predictions; normal acinar, metaplasia, and dysplasia predictions all override DAPI predictions.

### Validation and Testing

Because no foreign tissue was used for negative controls during training (primarily lymph nodes and GI tissue), regions of images containing these tissues had to be cropped out prior to testing and analysis. Testing and analysis were performed through a similar pipeline as training, incorporating intermediate crop normalization and tile level prediction. These overlapping tiles were stitched back into a full image and an average was taken to get pixel level predictions for each model. Model predictions were compared to immunostained serial sections that were thresholded by an expert. To do this, ADM and dysplasia predictions were combined to make a general pan-keratin prediction mask. Predictions were then paired with their respective serial section and correlated to determine model accuracy.

The amylase, pan-keratin and DAPI area were measured in pixels, and the percentage of positive areas were calculated as a percent of the total all measured cellular regions. The differences in means were assessed by independent-samples T-Test or one-way ANOVA. The correlation was tested by Bivariate Pearson analysis.

### Animal models

All animal use was approved by the OHSU Institutional Animal Care and Use Committee. The KC mice were all backcrossed at least 5 generations into the C57Bl6/J background. Acute pancreatitis was induced in 6-week old C57Bl6/J mice by intraperitoneal injection of 50 µg caerulein (Wisent INC:450-185-EG) per kg body weight, with a total of 7 consecutive treatments at 1hour intervals. Pancreatic tissues were harvested 3 days following caerulein treatment. Caerulein was dissolved in PBS at a concentration of 10 µg/ml.

